# Navigating a diversity of turbulent plumes is enhanced by sensing complementary temporal features of odor signals

**DOI:** 10.1101/2021.07.28.454116

**Authors:** Viraaj Jayaram, Nirag Kadakia, Thierry Emonet

**Author notes:** Contributed equally (Viraaj Jayaram, Nirag Kadakia).

## Abstract

We and others have shown that during odor plume navigation, walking *Drosophila melanogaster* bias their motion upwind in response to both the frequency of their encounters with the odor (Demir et al., 2020), and the intermittency of the odor signal, i.e. the fraction of time the signal is above a detection threshold (Alvarez-Salvado et al., 2018). Here we combine and simplify previous mathematical models that recapitulated these data to investigate the benefits of sensing both of these temporal features, and how these benefits depend on the spatiotemporal statistics of the odor plume. Through agent-based simulations, we find that navigators that only use frequency or intermittency perform well in some environments – achieving maximal performance when gains are near those inferred from experiment – but fail in others. Robust performance across diverse environments requires both temporal modalities. However, we also find a steep tradeoff when using both sensors simultaneously, suggesting a strong benefit to modulating how much each sensor is weighted, rather than using both in a fixed combination across plumes.

## Introduction

The complexity of natural odor plumes makes olfactory navigation a difficult task. Turbulent flows produce rapid changes in the local concentrations of the odor and instantaneous local gradients of the odor often do not point toward the source (Celani et al., 2014; Crimaldi and Koseff, 2001). Encounters between the animal and odorized packets of air are intermittent, with durations and frequencies spanning many orders of magnitude (Celani et al., 2014). Additionally, distinct flow conditions result in distinct spatiotemporal statistics: near boundaries and with lower mean wind speeds, odor plumes are smoother, with odor concentrations typically above detectable thresholds (Connor et al., 2018). But roughness in the physical landscape – sands, rough terrain, vegetation – and shifting winds can cause plumes to break up into discrete odor filaments, interspersed with long periods of undetectable concentrations (Carde and Willis, 2008; Murlis et al., 1992; Riffell et al., 2008).

To navigate plumes exhibiting this degree of temporal complexity, animals must be able to detect odor encounters quickly and accurately. Indeed, many organisms have evolved olfactory receptor neurons (ORNs) that respond to chemical signals with high temporal precision (Gorur-Shandilya et al., 2017; Jacob et al., 2017; Nagel and Wilson, 2011; Szyszka et al., 2014a; Szyszka et al., 2012). In *Drosophila melanogaster*, ORN firing responses are strongly time-locked to the arrival time of an odor (Gorur-Shandilya et al., 2017) and fast synaptic mechanisms (Fox and Nagel, 2021; Martelli et al., 2013) allow this information to be passed quickly downstream, within milliseconds, to projection neurons in the antennal lobe to drive rapid behavioral response (Bhandawat et al., 2010). Such precision has been suggested to allow accurate encoding of temporal features of the odor signal (Nagel et al., 2015), such as the frequency of odor arrivals.

In addition to these fast responses, *Drosophila* ORNs also adapt their firing rates and gain to prolonged stimuli (Cao et al., 2016; Gorur-Shandilya et al., 2017; Nagel and Wilson, 2011), priming them to accurately encode future odor signals (Kadakia and Emonet, 2019) without losing temporal precision as intensity changes (Gorur-Shandilya et al., 2017; Martelli et al., 2013). Likewise, in honeybees, temporal resolution of odor pulses increases with time in a pulsed odor environment (Szyszka et al., 2014b) and in the moth *Agrotis ipsilon*, ORN responses adjust in such a way as to optimally encode odor signals after blank periods of the likeliest duration (Levakova et al., 2018). Olfactory neurons in insects are thus sensitive to the temporal features of odor signals on both short and long timescales (Nagel et al., 2015).

Temporal precision in olfaction extends beyond insects. In mice, plume dynamics as fast as tens of milliseconds are encoded downstream in mitral and tufted cells (Ackels et al., 2021). In crustaceans, odors are encoded by bursting ORNs, or bORNs, which burst only if odors arrive at some phase relative to an intrinsic bursting cycle (Park et al., 2014). These cycles vary over orders of magnitudes across the bORN population, providing a natural template to encode the timing between odor arrivals (Park et al., 2016).

Naturally, such precisely-resolved temporal odor information shapes navigational decisions. When tracking pheromones, flying male moths fly faster and straighter upwind when receiving odor hits at higher frequencies than lower ones (Mafra-Neto and Carde, 1994; Vickers and Baker, 1994). Walking silkworm moths switch from zigzagging motion to straighter trajectories upwind in higher frequency environments (Kanzaki et al., 1992). In water, crabs navigate successfully in environments with higher odor intermittency, but fail to find odor sources as pulses become more infrequent (Keller and Weissburg, 2004). In straight ribbons, flying flies counterturn shortly after passing through the odor (Budick and Dickinson, 2006; van Breugel and Dickinson, 2014), and some models (Vickers and Baker, 1994) suggest that odor hits suppress an otherwise persistent internal counterturning mechanism, allowing moths to maintain straight trajectories if odors are frequent or long.

Two recent studies in *eLife* have quantified in great detail, using both experiment and extensive mathematical modeling, the olfactory navigational strategies of walking *Drosophila* in wind tunnels. One of these (Alvarez-Salvado et al., 2018) focused on spatially uniform but temporally varying environments, where the odor was presented in spatially uniform pulses lasting anywhere from 1 to 10 seconds. In this environment, walking flies maintained upwind headings and increased walking speed over the duration of the odor pulses, albeit with a degree of desensitization over time. This suggests that when odor encounters are long and persistent, the intermittency of the odor signal – the percentage of time the odor signal is above threshold – is a main driver of navigational decisions. The second study (Demir et al., 2020) instead challenged flies to navigate spatiotemporally complex odor plumes that were generated by stochastically perturbing a thin ribbon of odor. In this plume, odor encounters were much shorter (∼0.1-0.3s), more frequent (∼3 Hz), and less predictable. In that study, fly navigation was reproduced by a model in which only the frequency of odor encounters controlled upwind orientation, independent of their duration or concentration. These two studies used the same organism with the same locomotive repertoire. The two distinct models they uncovered naively suggests that flies are able to sense distinct temporal features of odor plumes, and use these various inputs to shape navigational decisions.

Here, we use mathematical modeling and numerical simulations to investigate how and under what conditions these two temporal features – odor intermittency and encounter frequency – can enhance the navigation of turbulent odor plumes. To examine the contribution to navigation from these two temporal features alone, we ignore all other sensory modalities, such as concentration sensing, concentration gradient sensing, bilateral sensing, and vision. We first demonstrate analytically that the dynamical model proposed in the first study above picks out (in appropriate limits) odor signal intermittency, while the model in the second study responds to the frequency of odor hits. These two temporal features are complementary and can be varied independently, forming a natural basis of temporal sensing. We devised a simple combined model that incorporates intermittency sensing and frequency sensing in a minimal way, and uses these two “sensors” to drive upwind orientations. Using agent-based simulations, we first show that this combined model requires both sensors to successfully navigate both measured plumes used in the two studies. We then applied the navigational model to simulated plumes, leveraging an advecting-diffusing packet framework that mimics odor motion in turbulent flows (Farrell et al., 2002). We find that to robustly navigate across a variety of plumes, agents should leverage both intermittency and frequency sensing. However, there is a steep tradeoff in performance when using both temporal features simultaneously, which persists across a variety of plumes. This predicts a strong benefit to modulating the weight of these two sensors, and we propose simple experiments to test whether flies or other insects indeed carry out such adaptation on slower timescales.

## Results

### Two experimentally-constrained models implicate distinct odor signal features in olfactory navigation

Our study is motivated by two models recently extracted from experimental observations of walking *Drosophila* navigating odor plumes (Alvarez-Salvado et al., 2018; Demir et al., 2020). Here we examine how they each respond to distinct temporal features of the odor signal. In the first model (Figure 1A) (Alvarez-Salvado et al., 2018), the instantaneous odor concentration *odor*(*t*) is first compressed into the range 0 to 1 using an adaptive Hill function

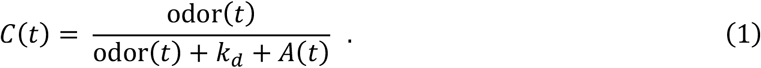

The half-max is set by *A*(*t*), a low-pass filtered sliding average of the instantaneous odor concentration

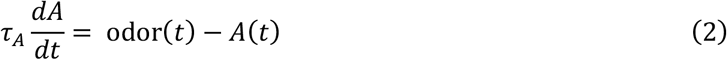

This mimics the gain adaptation of ORNs to the mean signal (Cao et al., 2016; Gorur-Shandilya et al., 2017). At the onset of a sudden increase in odor concentration, the compressed signal *C*(*t*) increases instantaneously before relaxing back to ∼0.5 with timescale *τ*_*A*_ = 9.8 sec. The compressed signal *C*(*t*) is then exponentially filtered into an “ON” function,

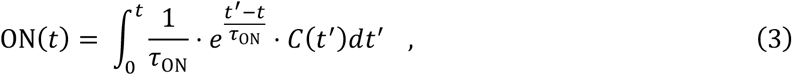

which drives odor-elicited behavioral actions. When ON(*t*) is high, the fly accelerates and biases its heading upwind; when ON(*t*) is low, the fly’s orientation randomizes and drifts downwind and its walking speed reduces (Alvarez-Salvado et al., 2018). We show analytically (below) that the value of ON(*t*) – and therefore the navigational actions – are largely determined by the intermittency of the odor signal, defined as the percentage of time an odor signal is present. Thus, we refer to this model as the *intermittency model*.

**Figure 1:**
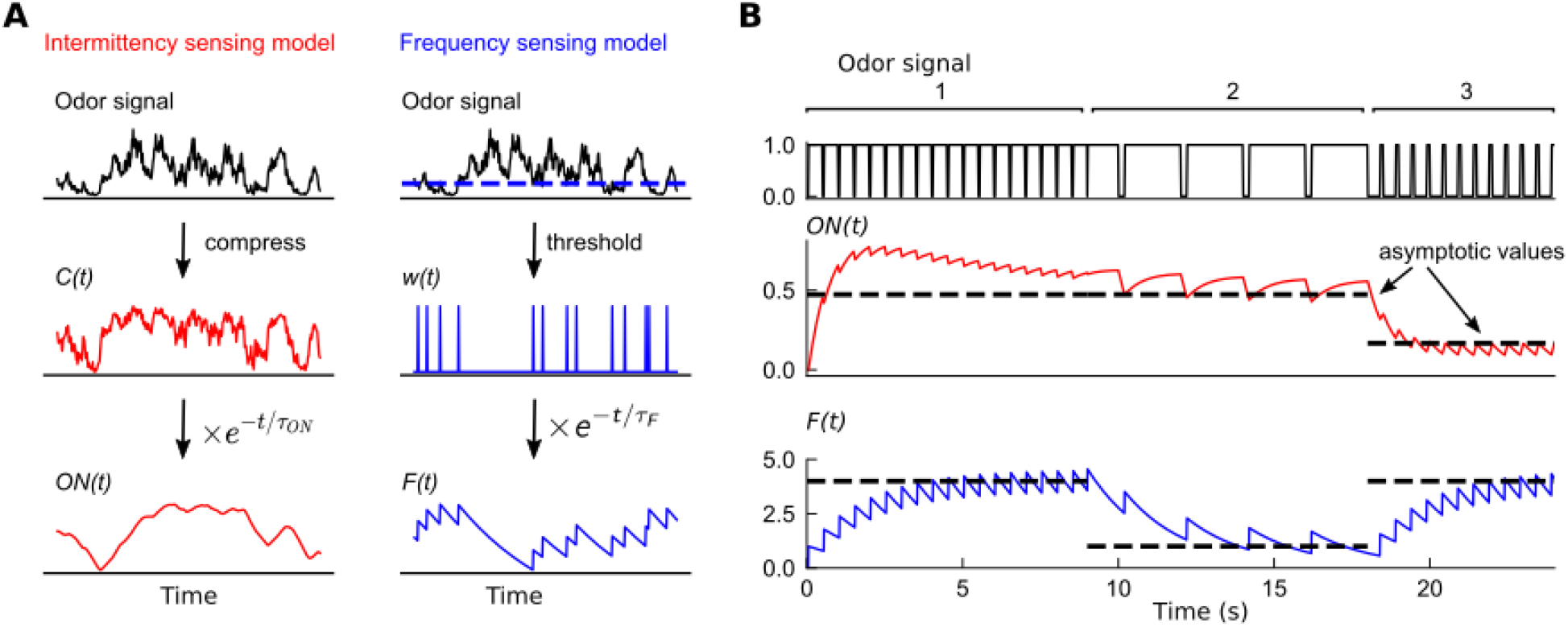
Filters extracted from experiment capture distinct temporal features of odor signals. **A**. Two experimentally-informed models (Alvarez-Salvado et al., 2018; Demir et al., 2020) of *Drosophila* olfactory navigation transform odor signals in distinct ways. Left column: the intermittency model compresses the odor signal with an adaptive nonlinearity into a representation *C*(*t*), bounded between 0 and 1. *C*(*t*) is then exponentially filtered with timescale *τ*_*ON*_ to generate *ON*(*t*). Right column: the frequency model thresholds the odor signal (dashed line in top plot) into a binary representation *w*(*t*), which is then passed through an exponential filter with timescale *τ*_*F*_ to generate *F*(*t*). **B**. Response of each of the models (bottom two plots) to a binary odor signal (top plot) of high intermittency, high frequency (Region 1), high intermittency, low frequency (Region 2) and low intermittency, high frequency (Region 3). The intermittency model is sensitive to the intermittency of the signal – in regions 1 and 2, it approaches a high value asymptotically, but a low value when intermittency is low, even if the frequency remains high (Region 3). The asymptotic values of the intermittency model (dashed lines) are 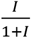 where *I* is signal intermittency (Methods for derivation). Conversely, the frequency model exhibits sensitivity to the frequency of encounters, tending asymptotically towards *fτ*_*F*_ where *f* is the signal frequency (dashed line). The frequencies in the three regions are 2Hz, 0.5Hz, and 2 Hz, the encounter durations are 0.45s, 1.8s, 0.1s, and the intermittencies are thus 0.9, 0.9, and 0.1.

In the second model (Figure 1A) (Demir et al., 2020), a detection threshold is used to detect when the odor arrives. This results in a binary time series *w*(*t*), which spikes as a *δ*-function each time the odor concentration crosses the threshold from below, and is 0 otherwise. The frequency of odor encounters is then estimated by filtering *w*(*t*) with an exponential:

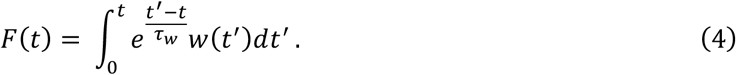

Thus *F*(*t*) rises by 1 at each threshold-crossing, before decaying exponentially with timescale *t*_*w*_ until the next odor hit. In this model, *F*(*t*) plays a similar role as ON(*t*) in the previous model in that it drives behavioral response to odors. When *F*(*t*) increases, flies increase their bias upwind and stop less and for a shorter time (Demir et al., 2020). Since *F*(*t*) is effectively a running average of the frequency of odor hits, we refer to this model as the *frequency model*.

To illustrate how each of these two sensory modalities respond to the temporal features of odor signals, we plotted the output of each filter in response to square-wave odor pulses of given frequency and intermittency (Figure 1B). These two features can be independently tuned – an odor signal can be high frequency and high intermittency if the whiffs (periods above threshold) are interrupted frequently with blank periods that are very short (region 1 in Figure 1B), while it can have high intermittency but low whiff frequency if whiffs are interrupted with short blank periods occurring more sparsely (region 2 in Figure 1B). In the first 2 regions of the signal, where intermittency is high, the response of the ON(*t*) model approaches a high value after an initial transient, while it drops to a lower steady state in region 3 where the signal intermittency is lower. The steady state response of ON(*t*) model is sensitive to the signal intermittency, but is independent of the whiff frequency, as indicated by the average response asymptote *I*/(1 + *I*), which monotonically increases with intermittency (Methods). In contrast, the frequency model responds strongly in regions 1 and 3 where whiff frequency is high, consistent with its asymptotic response *f* · *τ* (Methods). This happens irrespective of the disparity in signal intermittency between these regions (Figure 1B, bottom trace). Note that both models are sensitive to the temporal characteristics of the signal, but not absolute concentration.

Though these two models were extracted from the same model organism with the same locomotive repertoire – fruit files walking in a 2D arena – the experiments were performed in very different odor and flow conditions. The intermittency model was first extracted from flies navigating a uniformly odorized region of odor within a laminar airflow (Alvarez-Salvado et al., 2018). Using simulations, the model was then shown to qualitatively recapitulate navigational behavior in a measured near-bed turbulent plume (Connor et al., 2018) (Figure 2A), which we call the high intermittency plume, in which the odor signal was ever-present and varied on relatively long timescales of several seconds or more (Figure 2B). In contrast, the frequency model was fit to trajectories of flies navigating a plume with a high degree of spatial complexity (Figure 2C) generated by perturbing a fast laminar flow with stochastic lateral jets, which we call the high frequency plume. In that experiment, odor whiffs occurred frequently (2-5 Hz) (Figure 2E-F) and were much shorter (∼100ms) (Figure 2D). The two navigational models these experiments informed were clearly shaped by the plumes’ natural features: in the first, odor intermittency reached as high as 100% and whiff frequencies rarely surpassed 1 Hz (Figure 2E), whereas in the latter, the signal had intermittency mostly below 30% but whiff frequencies of several Hz (Figure 2F). Together, these two experiments and corresponding models suggest that flies use both odor frequency and intermittency to navigate upwind in different environments. This prompted us to ask how this dual sensing capability might enhance the efficacy and robustness of navigation in different conditions.

**Figure 2:**
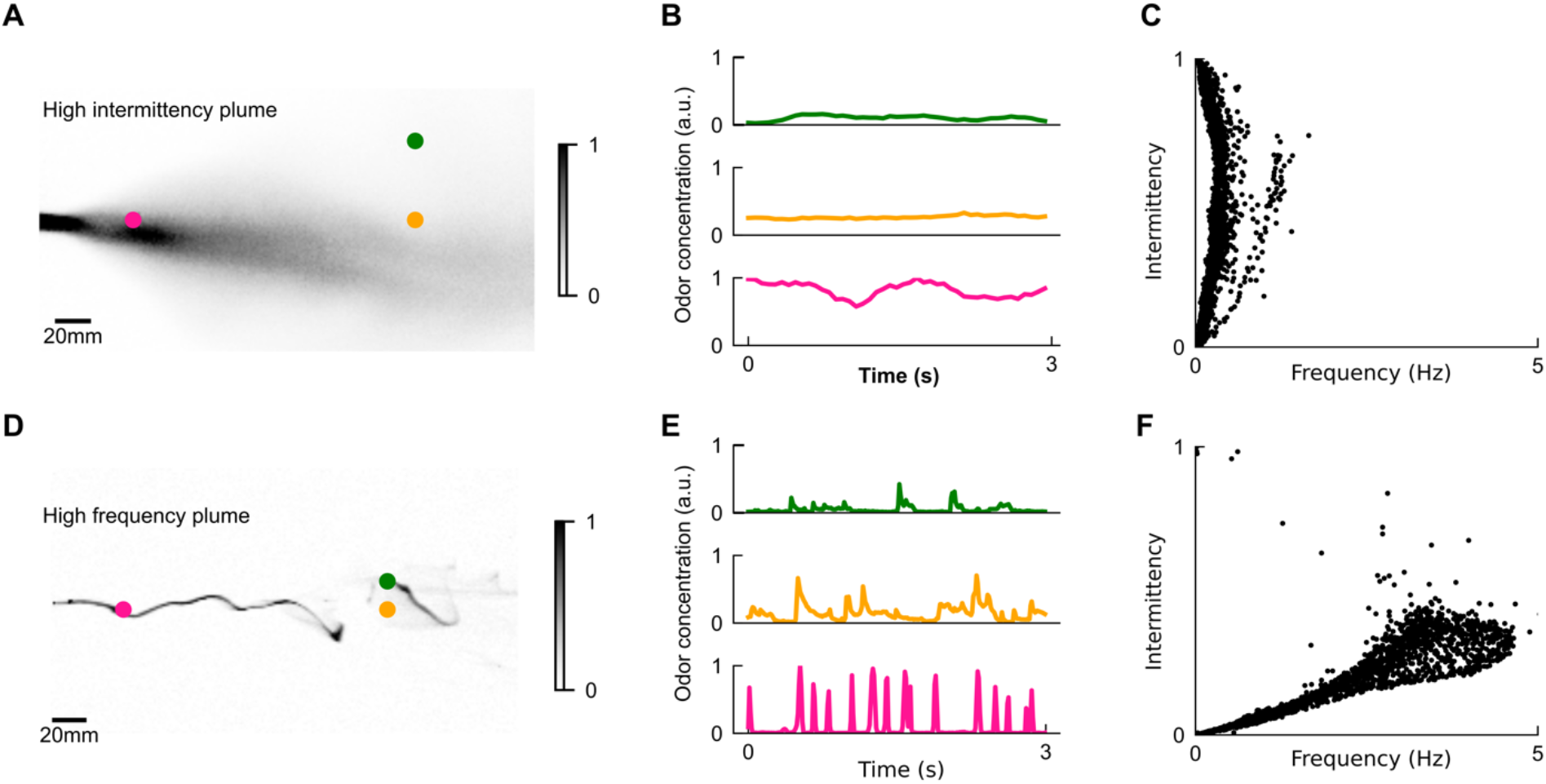
The differing temporal statistics of odor plumes. **A**. Snapshot of measured high intermittency plume, reproduced from data in (Connor et al., 2018). Colored dots: locations corresponding to odor series in in B. **B**. Odor concentration time series at different locations in high intermittency plume. **C**. Intermittency versus whiff frequency for 10,000 uniformly distributed points in the high intermittency plume. Statistics were calculated over the length of the full video. We see a range of intermittencies and many points with high intermittencies but relatively low frequencies. **D-E**. High frequency plume and representative time series, reproduced from data in (Demir et al., 2020). **F**.) Analogous to C for the high frequency plume. Data is clustered within a higher range of frequencies but low intermittencies.

### Dual intermittency and frequency sensing enhances navigation robustness in distinct environments

To address this question, we incorporated the two sensing capabilities, intermittency sensing and whiff frequency sensing, into a simpler model that eliminated some parameters. First, we simplified the *ON*(*t*) model into the following intermittency sensor

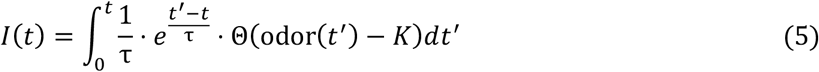

in which the adaptive compression from the original *ON*(*t*) sensor has been replaced with a thresholded odor signal. Θ is the Heaviside step function and *K* is the odor detection threshold. We kept the frequency sensor *F*(*t*) the same as above (Eq. 4), and used the same filtering timescales *τ* = *τ*_*ON*_ = *τ*_*w*_ for both the *I*(*t*) and *F*(*t*) sensors.

It is known that odor signals influence many behavioral actions, including speeding, turning, stopping (Alvarez-Salvado et al., 2018; Baker and Vickers, 1997; Demir et al., 2020; Mafra-Neto and Carde, 1994; Vickers and Baker, 1994). Given the near-universal response of insects to turn upwind or bias their turns upwind in the presence of odor (Baker et al., 2018), here we assumed agents walk at a constant speed unless they are turning, and focused on signal-driven changes in orientation. Turns occur randomly at a Poisson rate *λ*_*turn*_ and turn magnitudes are sampled from a normal distribution 𝒩(30^°,^ 8°) as found before (Demir et al., 2020). Turn directions (sign of the orientation change) are modeled as:

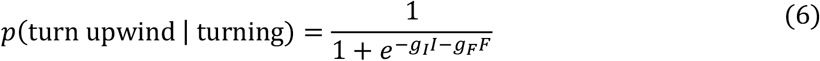

Thus, the likelihood that a turn is directed upwind (versus downwind) increases sigmoidally with a linear combination of *F*(*t*) and *I*(*t*). In the absence of signal, upwind and downwind turns are equally likely: *P*(upwind|turn) = 0.5 (further specifics of parameters choices in Methods). For now, the “sensor gains” *g*_*I*_ and *g*_*F*_ were set to 1.9 and 0.2, respectively, by comparing to experimental data (Methods). We refer to these values as the “base gains” and denote them as *g*_*I*0_ = 1.9 and *g*_*F*0_ = 0.2. We also define an intermittency-only and frequency-only sensing model by setting *g*_*F*_ and *g*_*I*_ to 0, respectively.

To examine how frequency and intermittency contribute to navigational performance in this combined model, we simulated *N* agents navigating both the high intermittency and high frequency plumes. The initial position and orientations of the agents were randomized uniformly. Performance was quantified as the fraction of agents that reach within 15 mm of the source in the presence of an odor signal,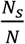, minus the fraction of agents,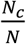, that reach the source by chance, i.e. when no signal is present. Individual trajectories of successful flies in either plume look similar: when oriented away from the source, agents are quickly able to reorient within the plume region, and navigate to the source with relatively straight trajectories combined with occasional corrective kinks (Figure 3B). Overall, agents navigated successfully in both plumes (Figure 3C), and performance was relatively robust to initial angle and position (Figure 3D). However, when either frequency sensing (*g*_*F*_ = 0) or intermittency sensing (*g*_*I*_ = 0), were removed, performance degraded (Figure 3D) in one of the plumes, and became more sensitive to initial conditions. Though not wholly surprising that removing sensors degrades performance, this suggests that a simple linear combination robustly navigates two disparate odor plumes, without exhibiting any obvious failure modes due to interference between sensors.

**Figure 3:**
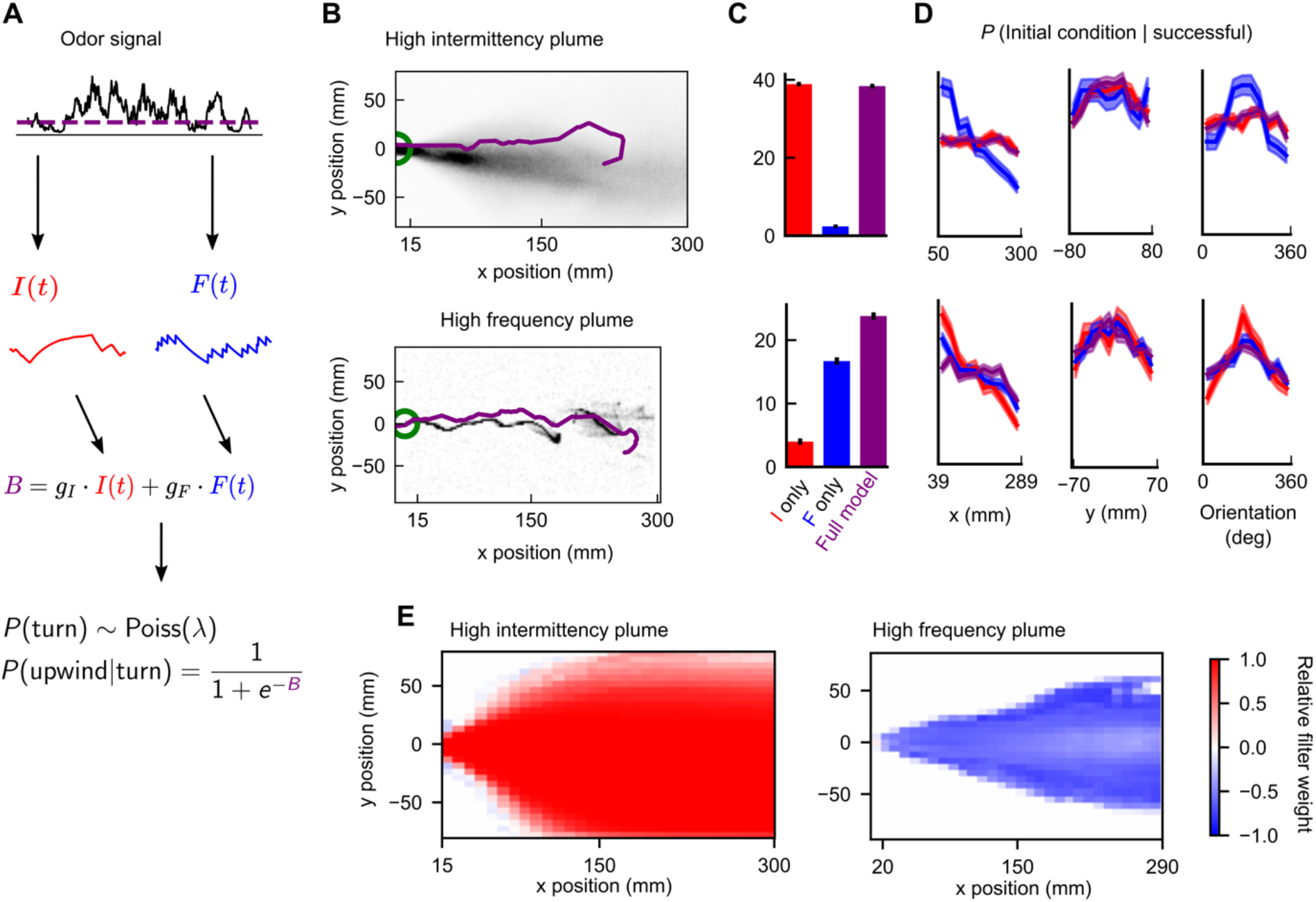
Sensing both intermittency and frequency enables navigation across diverse plumes. **A**. Our model linearly combines an intermittency sensor (red) and whiff frequency sensor (blue) to bias upwind motion. Following (Demir et al., 2020), turns occur stochastically at a constant Poisson rate *λ*_*turn*_, while the sensor output *B* biases the likelihood that turns are upwind. Turn magnitudes are chosen from a normal distribution with mean 30° and S.D. 8° (Demir et al., 2020). **B**. Example successful trajectories in the high intermittency and high frequency plume (Figure 2). **C**. Percentage of agents that reach within 15 mm of the source when signal is present minus same percentage when signal is absent, for the model with only intermittency sensing (*g*_*F*_ *=* 0; red), only frequency sensing (*g*_*I*_ *=* 0; blue), or both (*g*_*F*_, *g*_*I*_ nonzero; purple), in the high intermittency plume (top) and high frequency plume (bottom). Error bars: SEM calculated by bootstrapping the data 1000 times (Methods). **D**. Distribution of initial downwind position *x* (first column), crosswind position *y* (2^nd^ column) and orientation (3^rd^ column) for successful agents, for the high intermittency (top row) and high frequency (bottom row) plumes. Colors correspond to same models as in D. Upwind heading is 180° and shaded regions represent SEMs obtained from bootstrapping (Methods) **E**. Relative filter 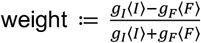 for different points in the two plumes.

We expect that the two sensors do not contribute equally at all times to the navigation and that the relative contribution of either sensor may depend on location within the plume – for example, in the high frequency plume, the intermittency sensor might be more active near the plume centerline, where the signal is more likely to be present. To quantify this, we measured the relative weight of each sensor 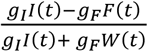, which interpolates between pure intermittency sensing (+1) and pure frequency sensing (−1). As expected, the intermittency sensor dominates in the high intermittency plume, whereas the frequency sensor dominates in the high frequency plume (Figure 3E). Still, this dominance is not absolute. For example, frequency sensing plays a role near the conical boundary of the high intermittency plume. Likewise, intermittency contributes significantly along the centerline of the high frequency plume. These modest but significant contributions led us to next wonder how the sensors might be relatively weighted to optimize navigational performance, and how this weighting might change in different plumes.

### Optimal performance requires distinct weighting of frequency and intermittency in different environments

To investigate the influence of relative sensor weight in navigation, we quantified navigational performance as a function of both the sensor weights *g*_*I*_ and *g*_*F*_ and the plume’s spatiotemporal complexity. To remove constraints due to the limited spatial and temporal resolution of the recorded plume videos, and to easily investigate a wide range of environments, we switched to simulated odor plumes. The plumes were modeled using a simple dispersion model (Farrell et al., 2002). Gaussian packets of odor are released from a source at a fixed Poisson rate *λ*, and advected by a velocity field composed of a uniform downwind velocity *U*. Normally distributed random perturbations *η*_*x*_ and *η*_*y*_ are added to the packet positions in the crosswind and downwind directions, respectively, at each time step, to account for the effects of turbulent diffusivity. The turbulent diffusivity models the effects of turbulent eddies as a diffusive process, but with diffusion constant κ that can greatly exceed molecular diffusivity. In addition, the Gaussian packets grow in size with an effective diffusivity *D*, to account for the combined effects of molecular diffusion and smaller eddies in the wind flow (Figure 4A-B). Varying *U* and *D* allowed us to generate plumes with diverse temporal statistics. *U* 2 36mm/s and *D* 2 *F*2mm^2^/s resulted in a high intermittency plume with long whiff duration (Figure 4C). Increasing the wind speed to *U* 2 300mm/s and decreasing effective diffusivity to *D* 2 10mm^2^/s resulted instead in a higher frequency plume with much shorter whiffs (Figure 4D). In each plume, we simulated 10,000 agents with uniformly distributed initial position and heading angle, where each agent navigated with a fixed set of gains *g*_*I*_ and *g*_*F*_. We investigated various choices of *g*_*I*_ and *g*_*F*_, from 0 to 50X the base gains (Methods).

**Figure 4:**
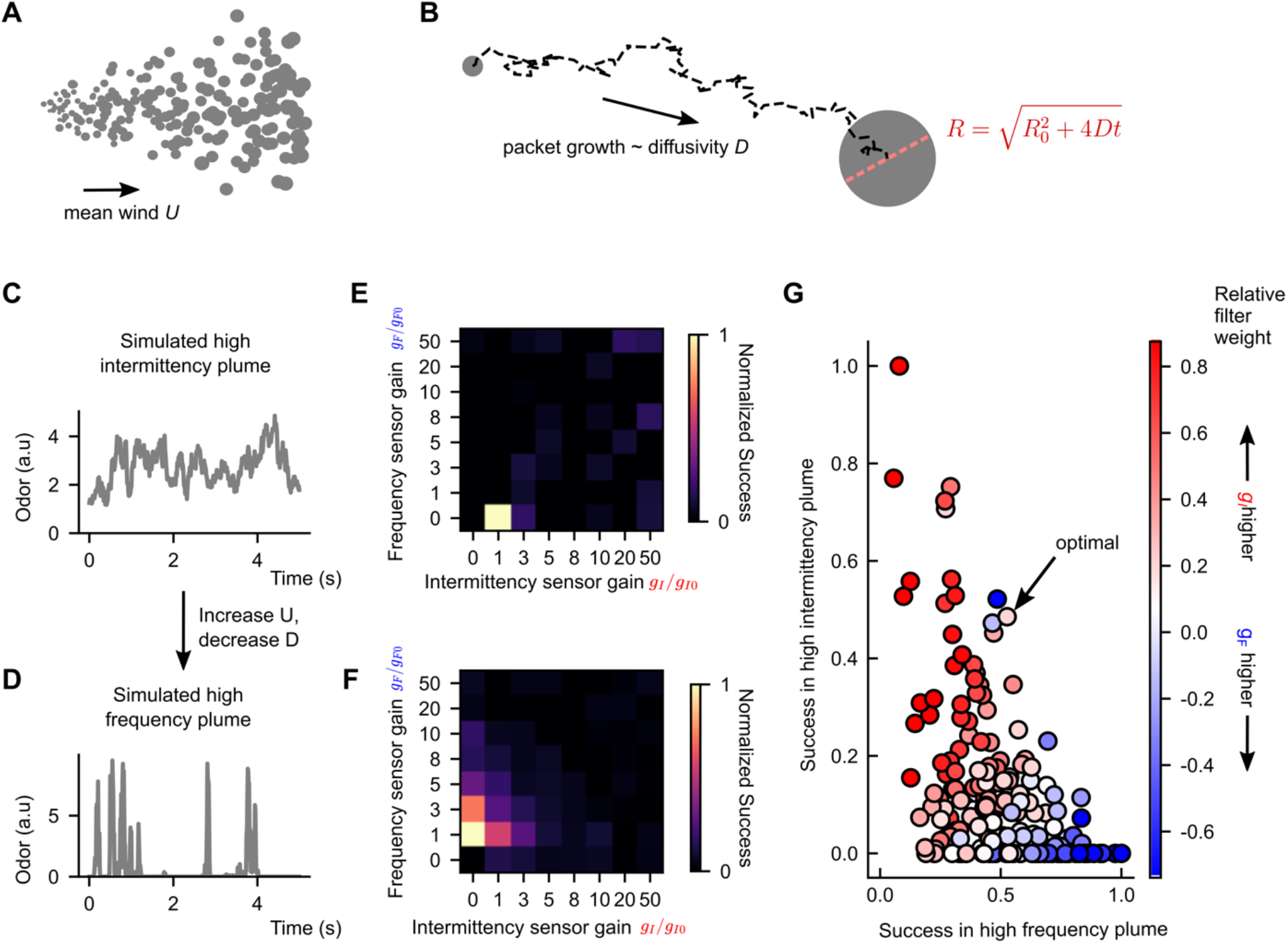
Performance tradeoff between intermittency and frequency sensing in two diverse turbulent plumes. **(A)** Example of a simulated odor plume, following the framework in (Farrell et al., 2002). Gray circles denote Gaussian odor packets. **(B)** Example trajectory of a single odor packet in these simulations, and illustration of its growth. **(C)** Example odor concentration time series in a simulated high intermittency plume. **(D)** Same as C, for a high frequency plume. **(E)** Normalized success percentage within the simulated high intermittency plume (where success percentage is calculated as in Figure 3C) for different sets of *I* and *F* gains, after adding noise to *I* and *F*. Gains are measured in multiples of the base gains, defined in Methods. **(F)** Same as E, but for the simulated high frequency plume. **(G)** Normalized success percentage in the high intermittency plume versus same in the high frequency plume; each dot represents a different set of gains *g*_*I*_ and *g*_*F*_. Points are colored by the relative weighting of the two sensors (see Methods for calculation details). Indicated point: gains that maximized the geometric mean of normalized success across the two plumes. The concavity of the front suggests a tradeoff in performance in either plume.

The *g*_*I*_ and *g*_*F*_ maximizing performance in our simulated high intermittency plume was reasonably constrained, with a clear maximum occurring around the experimentally motivated gain (Figure 4 Supplement). However, in the simulated high frequency plume, a variety of gains led to similarly maximal performance (Figure 4 Supplement), including some with values an order of magnitude larger than the base gains. We suspected that while a heavy weighting of either sensor might improve navigation, such extreme amplification may also compound the effects of noise, leading to a lack of robustness in natural conditions. We therefore added Gaussian noise to the *I* and *F* filters – noise amplitude was 5% of the average value of *I* (F) in the center of the simulated high intermittency (high frequency) plume. This removed maxima at high gains but retained clear maxima at lower gains (Figure 4E-F). Interestingly, the unique maxima sat fairly close to the base gain values (e.g. values of 1 in Figure 4E-F). In addition, the optimal gains for the simulated high intermittency and high frequency plumes had *g*_*F*_ = 0 and *g*_*I*_ = 0, respectively. This inherent trade-off illustrates that simply augmenting the sensory capability can at times degrade performance, suggesting a benefit for sensor specialization in distinct environments.

### Performance tradeoff between intermittency-sensing and frequency-sensing in different environments

To get a better understanding of how navigational performance in these two simulated plumes depends on the sensor weights, we did a tighter sweep of gains near the performance maxima (Figure 3E-F) for each plume. For each set of gains, we then plotted performance in the high intermittency plume against that in the high frequency plume. For comparison, we also plotted the set of gains 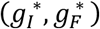 that maximized the geometric mean of normalized success in both plumes (indicated in Figure 4G). The resulting scatterplot quantifies the performance in the two plumes for different navigational models, where each model is parameterized by its sensor weights *g*_*I*_ and *g*_*F*_. In general, the scatter plot fills out a region near the origin, bounded by a curve that forms a “Pareto front” of navigational performance. This Pareto front reveals a performance tradeoff for the different models: combinations of *g*_*I*_ and *g*_*F*_ that are weighted toward *I* do better in the high intermittency plume, while combinations weighted toward *F* outperform in the high frequency plume (Figure 4G). There was no fixed set of gains that performs optimally in both plumes. Importantly, the apparent concavity of the Pareto front illustrates a somewhat steep tradeoff, and suggests that flies might be better off modulating gains and switching between using intermittency and frequency sensors to bias upwind motion, as opposed to using both simultaneously.

Finally, we wondered how this tradeoff manifests across a more diverse spectrum of plumes. The computational simplicity of the turbulent plume model allowed us to study a wide array of turbulent plumes differing in their temporal statistics. We fixed the gains to the values that optimized the geometric mean between the high intermittency and high frequency plumes, 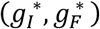 and then varied the environmental parameters *U* and *D* to smoothly interpolate between the high frequency and high intermittency plumes investigated above. Success was roughly uniform in the different environments (Figure 5A). However, removing the frequency sensor (*g*_*F*_ 2 0) significantly improved performance in the slowly advecting and highly diffusive plumes (low *U*; high *D*), which tend to be smoother in their concentration profiles. The reverse was true when we removed intermittency sensing (*g*_*I*_ 2 0), exemplifying a tradeoff in navigational performance that persists across this wide range of odor environments. Together with the results presented above (Figure 3), this suggests that while a naïve summation of temporal sensors may be beneficial in some cases, in general, navigation can always be improved by some degree of specialization.

**Figure 5:**
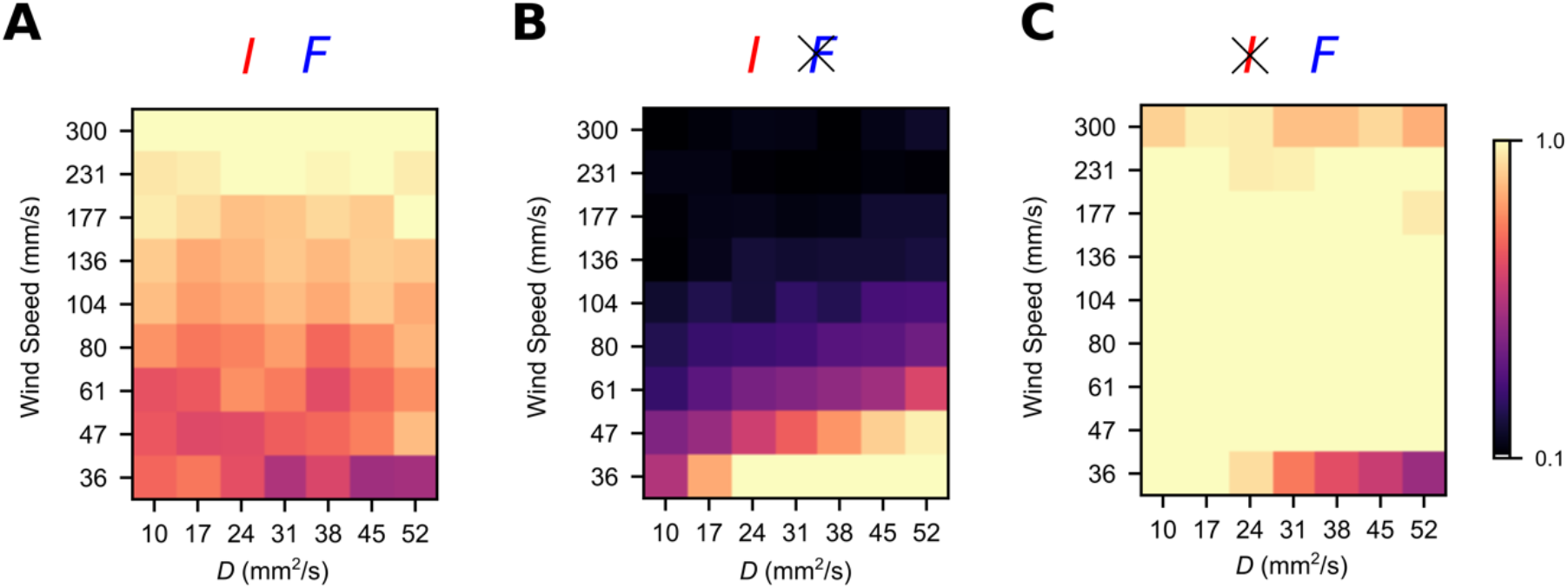
Simultaneous intermittency and frequency sensing maintains steady performance across a spectrum of odor environments, but does not allow for optimal performance. Normalized success percentage for a frequency and intermittency-sensing model (**A**), only intermittency-sensing model (**B**) and only frequency-sensing model (**C**) for a range of simulated odor plumes. Success percentage is normalized such that the best performance of the three models is set to 1 for each environment. Gains for (**A**) were chosen to optimize the geometric mean of performance in the simulated high intermittency and high frequency plumes. Gains in (**B**) and (**C**) were chosen by taking the gains in (**A**) and then setting *g*_*F*_ (**A**) and *g*_*I*_ (**C)** to 0.

## Discussion

In this work, we used numerical simulations to explore the value of two temporal features of the signal – odor intermittency and encounter frequency – in navigating naturalistic odor plumes spanning a range of spatial and temporal complexity. These two features are a natural set in that they can be varied independently to create a variety of odor signals (Figure 1). Other complementary and complete quantities could be used, such as whiff and blank duration (Rigolli et al., 2021), but we focused on these since they are directly implicated by various experiments in walking *Drosophila melanogaster*. The navigation model we proposed reduces two experimentally-informed models of fly olfactory navigation into elementary transformations that separately extract odor intermittency and encounter frequency, and then uses these two “sensors” to bias the agent upwind. An interesting finding here is that the optimal agent in the two simulated plumes assigned weights to the sensors that resembled the weights inferred from experiment (Demir et al., 2020) (Figure 4E, Methods). Loosely, this suggests that the manner in which temporal features are extracted and processed within the *Drosophila* olfactory circuit may already be adapted to natural plume environments.

We emphasize that our work explores normative strategies, so our results have no bearing on whether such adaptation actually occurs. There is, however, evidence that such adaptation may exist at the level of individual neurons: for example moth ORNs adjust their encoding efficiency to the local statistics of pheromones (Levakova et al., 2018). Additionally, we emphasize that in (Demir et al., 2020), upwind orientation was found to be independent of intermittency for fixed frequencies, suggesting that such adaptation of sensor weight may actually be present in walking *Drosophila*. Our work suggests future experiments, based on simple modifications of existing experimental paradigms, that could be used to quantify this slower-scale adaptation. One could present the complex odor plumes we generated in our recent work (Demir et al., 2020), while modulating the overall statistics on a slower scale via the speed or strength of the upwind lateral perturbations, the wind speed, or both, and record how upwind orientation depends on frequency or intermittency.

In the latter half of this study, we simulated odor plumes using a simple drift-diffusion model (Farrell et al., 2002) to efficiently generate a diversity of plumes. A more precise approach would be to numerically integrate the Navier-Stokes equations describing the wind flow, together with advective-diffusive scalar transport describing the dispersion of a scalar concentration field (Rigolli et al., 2021). In such simulations, resolving odor concentrations to the viscous scale is very computationally expensive. This would likely preclude the investigation over more than a handful of distinct odor plumes, as our simplified model allowed us to explore here. On the other hand, such detailed simulations show that even in a single plume, the statistics of the odor change significantly with distance from the source, and therefore animals may benefit from modulating sensory strategies during navigation (Rigolli et al., 2021). This is consistent with our finding that frequency sensing contributes more near the edges of the plume than it does near the centerline, and vice versa for intermittency sensing.

There are several aspects of olfactory navigation not considered in this work. In particular, we have neglected the role of bilateral sensing between the two antennae. In insects, bilaterally-resolved concentration sensing has been demonstrated in flies (Gaudry et al., 2013) and implicated in navigation of laminar ribbons (Duistermars et al., 2009). Bilateral sensing has also been demonstrated in mice (Rajan et al., 2006), sharks (Gardiner and Atema, 2010), and even humans (Wu et al., 2020), and has been implicated in effective navigation in aquatic environments (Michaelis et al., 2020). Spatially resolved information has been shown theoretically to provide more information about an agent’s position relative to the source of the odor (Boie et al., 2018) and aid olfactory navigation strategies, even in plumes with elements of stochasticity and turbulence (Hengenius et al., 2021). For very closely spaced antennae as in flies (<1 mm), these gradients are very difficult to resolve and so are often not useful for navigation (Celani et al., 2014; Crimaldi and Koseff, 2001; Shraiman and Siggia, 2000). Nonetheless, it would be interesting to consider the effect of bilateral comparisons of intermittency and frequency, particularly when modeling the navigation of species with larger antennae.

To this end, it has already been shown that bilateral comparisons of frequency allow agents to track the edges of some turbulent odor plumes (Michaelis et al., 2020). Additionally, recent work (Rigolli et al., 2021) has shown that in the central regions of a turbulent plume, the intensity of an odor signal is more useful than temporal statistics for predicting distance to source, while at the edges of the plume, temporal statistics become more useful. Thus it is very possible that in high intermittency plumes, organisms might use frequency to track the edges of odor plumes or even execute offset responses, such as those detailed in (Alvarez-Salvado et al., 2018).

For the sake of simplicity, we considered a model where agents move with a constant speed and only change orientation through a discretized turning paradigm, suggested by (Demir et al., 2020). However, more diverse actions such as stopping and walking (Demir et al., 2020), speed modulation (Alvarez-Salvado et al., 2018; Mafra-Neto and Carde, 1994), continuous heading modulation (Alvarez-Salvado et al., 2018) and casting/counter-turning behavior (Alvarez-Salvado et al., 2018; Budick and Dickinson, 2006; Mafra-Neto and Carde, 1994; Pang et al., 2018; Vickers and Baker, 1994) have also been observed in insect olfactory navigation. It is also worth investigating, therefore, the role of intermittency and frequency in modulating behaviors such as these in different environments.

Furthermore, due to the thresholding and compression of the odor signal in the two experimentally-informed models that motivated this study, we have not investigated the role that odor concentration may play in modulating navigational behavior. Absolute odor concentration can inform source location in turbulence (Rigolli et al., 2021), and more information is conveyed about spatial location when resources are devoted to encoding higher absolute concentrations (Victor et al., 2019). Concentration sensing has also been implicated in neuron response: in moth projection neurons, spike patterns depend on both the intensity and timing of odor stimuli (Vickers et al., 2001). Thus, it is likely that navigational performance could be enhanced by incorporating more information about the intensity of the odor signal, along with the extracted temporal statistics.

Finally, we have not explored the role of learning. The frequency and intermittency filters we used had a timescale of two seconds, precluding history-dependent behavioral effects over longer timescales. History-dependence in navigational decisions has been observed in flying fruit flies (Pang et al., 2018), where the magnitude of fly turns decreased with the number of signal encounters, in desert ants (Buehlmann et al., 2015), where ants used the existence of previously learned olfactory cues to navigate in a new environment, and in mice (Gire et al., 2016), where gradient climbing was abandoned for foraging when mice were sufficiently conditioned on known odor locations. Theoretical strategies such as infotaxis, where agents navigate by using cues to learn an internal probabilistic representation of their environment (Vergassola et al., 2007), also has some support in experiment (Calhoun et al., 2014; Pang et al., 2018). We find that robust navigation is enhanced by modulating intermittency and frequency sensing in time, and incorporating history-dependence in our models could be done straightforwardly, with a few added parameters. Pairing this with behavioral experiments of the type suggested above would provide a fruitful direction for future study.

## Acknowledgements

We thank Mahmut Demir for providing measured plume data and advising on the simulations, Hope Anderson for help with agent-based simulation codes, and Henry Mattingly and Aarti Sehdev for helpful discussions. VJ and TE were supported by the Program in Physics, Engineering and Biology at Yale University. NK was supported by a postdoctoral fellowship through the Swartz Foundation for Theoretical Neuroscience, by postdoctoral fellowship NIH F32MH118700, and by postdoctoral fellowship NIH K99DC019397.

## Competing interests

The authors declare that no competing interests exist.

## Methods

### Simulating ON and F Responses to square waves

The frequency response function is defined as the convolution between the whiff onset time series *w*(*t*) and an exponential filter with decay timescale *τ*_*F*_ where the whiff time series is a sum of delta functions occurring at the onset of each whiff. Thus, we have

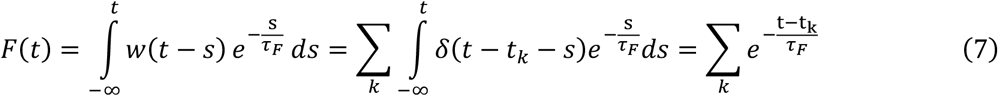

where *k* enumerates the whiffs. Note that 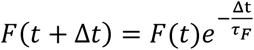. Therefore, in discrete time steps we have *w*(*t* + Δ*t*) = 1 if *odor*(*t*) < *K* and *odor*(*t* + Δ*t*) ≥ *K* and 0 otherwise and 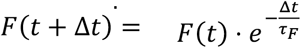, if *w*(*t* + Δ*t*) = 0 and 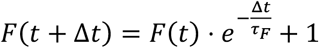 if *w*(*t* + Δ*t*) = 1.

For *ON*(*t*), we use Euler’s method to numerically integrate Equation (2) to obtain *A*(*t*) and then similarly integrate the following equation:

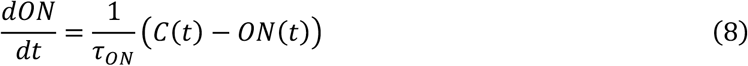

where *C*(*t*) is defined in Equation (1) and the above equation is equivalent to Equation (3).*τ*_*F*_ was set to 2s (Demir et al., 2020) while *τ*_*A*_ and *τ*_*ON*_ were set to 9.8s and 0.72s respectively (Alvarez-Salvado et al., 2018). The detection threshold was assumed to be below the signal amplitude and *k*_*d*_ was set to be 1% of the signal amplitude.

### Calculation of ON and F responses to square waves

To illustrate how the ON and W filters respond to the frequency and duration of odor signals, we consider their response to square wave odor pulses of given frequency *f*, duration *D* and amplitude *S*_0_. We first consider the ON response. To understand the ON response we first have to calculate *A*(*t*). From equation (2), we have

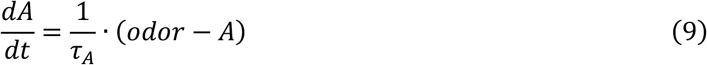

Let *A*_*n*_ denote the value of *A* at the *offset* of the *n*^*th*^ pulse of signal and 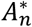 denote the value of *A* at the *onset* of the *n*^*th*^ pulse. We wish to obtain a recursive relation for *A*_*n*_ which will allow us to solve for *A*_*n*_ and from there obtain the value of *A* at all times. At the offset of a pulse, *odor* = 0 and *A* will exponentially decay with time scale *τ*_*A*_ until the onset of the next pulse. This time of decay is given by 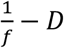. Hence at the onset of the next pulse, 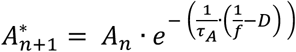. At this point, for a time period *D*, i.e. until the offset of the (*n* + 1)^*th*^ pulse, *A* obeys the equation

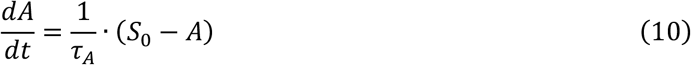

with initial value 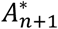 . Hence

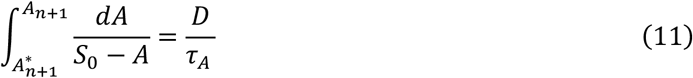

and therefore, after substituting 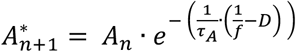

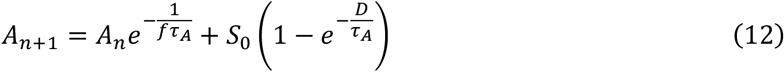

One can thus see that

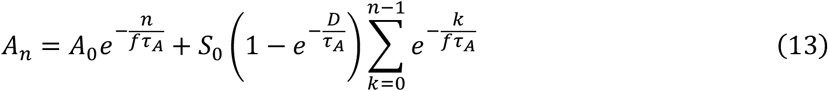

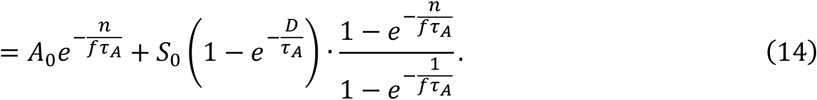

Once the number of pulses *n* is much greater than *fτ*_*A*_, i.e *t* ≫ *τ*_*A*_, we get

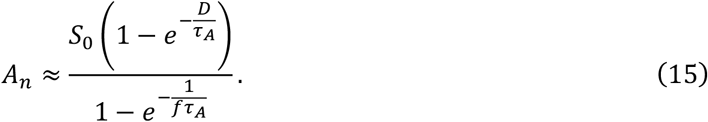

Since this is the value of *A*(*t*) at the end of a pulse, it will be the maximum value of *A*(*t*) over one period. Ultimately, however, we are interested in computing *ON*(*t*), which obeys the equation

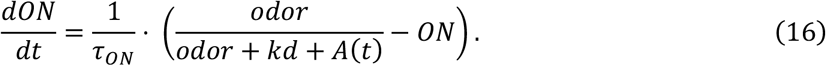

To understand the response of *ON* we can consider three different signal time scales. If the signal fluctuates quickly with respect to *τ*_*A*_ i.e. *D* and ^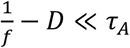^, then for *t* ≫ *τ*_*A*_ one can approximate *A*(*t*) with its average value over one period, which is given by

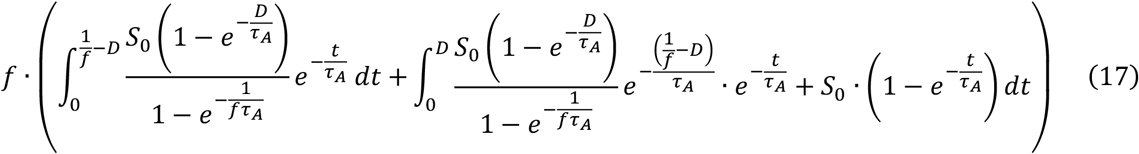

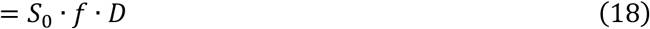

Notice *f* · *D* = *I*, the intermittency of the signal. Hence in this limit, and assuming *S*_0_ ≫ *kd*, when the signal is present we have

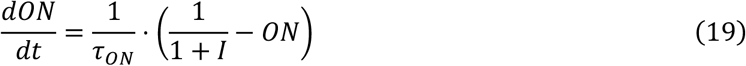

Thus, *ON*(*t*) obeys the same dynamics as *A*(*t*), except it adapts to a square wave of amplitude 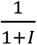 instead of *S*_0_ and with a different time scale. Thus by the same reasoning as for *A*(*t*), the maximum value of *ON*(*t*) over one period (once *t* ≫ *τ*_*A*_, *τ*_*ON*_) is approximately 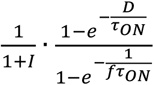 and the average value over one period is 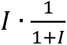 .

If instead *τ*_*A*_ ≈ *D* or *τ*_*A*_ ≪ *D* then *A*(*t*) ≈ *odor*(*t*) and we get

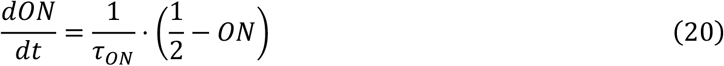

and the average value of *ON*(*t*) becomes *I*/2. (The maximum value would be 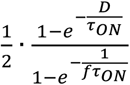).

Finally, we can consider the case where *τ*_*A*_ ≫ *D* and 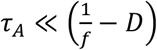. In this case *A*(*t*) ≈ 0 and *ON*(*t*) adapts to a square wave with amplitude ≈ 1. The average value of *ON*(*t*) is *I* (and the maximum value would be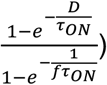).

In summary we see that in all these cases, the average value of *ON* depends only on the intermittency and increases monotonically with intermittency.

For *F*, it is easiest to consider *F*_*n*_ as the value of *F* just after the *onset* of each pulse. Since *F* increases by 1 at the onset of each pulse and then decays exponentially with time scale *τ*_*F*_ until the onset of the next pulse, one has

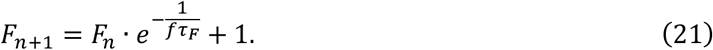

Hence

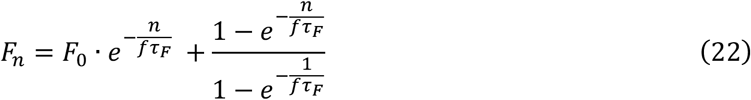

For *t* ≫ *τ*_*F*_ we have *n* ≫ *fτ*_*F*_ and 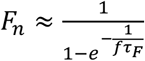. Since *F* jumps at the onset of a pulse and then decays, this is the maximum value of *F*. The average value of *F* over one period is thus

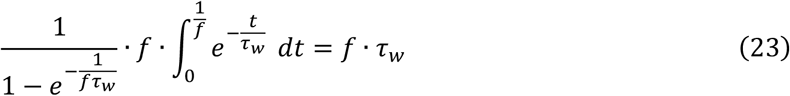

Hence the average value of *F* is linearly proportional to the frequency of the signal.

### Agent-based simulation in recorded odor plumes

The first plume recording we used is the same as used in (Alvarez-Salvado et al., 2018). We call this plume the high intermittency plume. The odor detection threshold of the agents was set by analyzing the signal in a region outside the plume. In this region, pixel values of 0 were removed and non-zero values were fit to a Gaussian. The detection threshold was then set to be the three standard deviations above the mean of this fit. 10,000 agents were initialized with uniformly distributed starting position, where the x-position was between 50mm and 300mm from the source and the y position went from 80mm below the source to 80mm above the source. The initial heading angle was uniformly distributed from 0 to 360 degrees. The simulation was run for the length of the video, (240s) and the discrete time step was set to be the reciprocal of the frame rate (1/15s).

The second plume recording we used was taken from (Demir et al., 2020). We call this the high frequency plume. The odor detection threshold of each agent was set the same way it was in (Demir et al., 2020). Again 10,000 agents were initialized with uniformly distributed initial position and heading. The initial x-position was between 38.45mm and 288.45mm and the initial y position was between -74mm and 86mm. Initial heading was uniformly distributed from 0 to 360 degrees. The simulation was run for 123.3s, starting from the 600^th^ frame of the video to the last frame, at 89.94 frames per second, corresponding to the frame rate used in (Demir et al., 2020). The first 600 frames were dropped so that the plume had expanded to full size when the simulations began.

In both simulations, odor signal was computed by averaging over an elliptical antenna-sensing region in front of the agent, as in (Demir et al., 2020). The length of the region’s major axis was 1.5mm and the length of the minor axis was 0.5mm. The ellipse was centered 1mm in front of the agent. In both simulations, if agents went outside the frame region then they were allowed to continue but received zero signal in those regions. Thus there were no walls in these simulations.

For these simulations, *F* was computed as for the square-wave pulses, with a detection threshold as described above, but we also enforced that the whiff time series *w*(*t*) could not register two whiffs less than 40ms apart, to capture the idea that the time resolution of individual whiffs is not arbitrarily precise and to avoid spurious detections due to the random fluctuations in the signal, as suggested by (Demir et al., 2020).

### Determination of base gains from experiment

The base gains, *g*_*I*0_ and *g*_*F*0_, which were used for the simulations in Figure 3, and in multiples of which the gains in Figure 4 and Figure 5 are reported, were determined the following way. (Demir et al., 2020) experimentally extracted a sigmoidal turning bias, as in Equation 6 except only using the *F* filter and reported a gain of 0.242. We thus set *g*_*F*0_ = 0.242. *g*_*I*0_ was set so that the contribution from *I* in the high intermittency plume would be roughly the same size as the contribution from *F* in the high frequency plume. So defining *I*_0_ and *F*_0_ to be typical *I* and *F* values in the high intermittency and high frequency plumes respectively, we have *g*_*I*0_*I*_0_ = *g*_*F*0_*F*_0_. We thus determined a *g*_*I*0_ of 1.936.

Regarding the remaining parameters, the turn-rate was set to 1.3/s, walking speed set to 10.1 mm/s, and filter decay timescale *τ* was set to 2s, all in accordance with the findings of (Demir et al., 2020). Note that the same timescale was used for the *I* and *F* filters.

### Statistical methods

Error bars for success rates (Figure 3C) were computed by bootstrapping data from a simulation of 10,000 flies-1000 resamples were used with each resample size being equal to 10,000. Similarly, for the histograms of successful initial conditions, the data was resampled 1000 times, where each resample size was the size of the original data and means and standard deviations were computed and used for each histogram bin.

### Agent-based simulation in simulated odor plumes

The simulated odor plumes were created using the strategy laid out by (Farrell et al., 2002). Plumes consisted of growing Gaussian packets of odor concentration, released as a Poisson process with rate *λ*, that were advected by a uniform mean wind velocity and perturbed by turbulent diffusivity. The concentration at a point (*x, y*) due to a packet centered at (*x*_*i*_, *y*_*i*_) was computed as

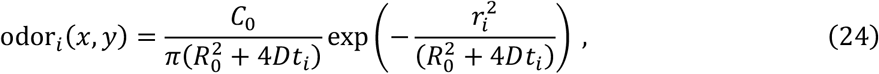

where *r*^2^ = (*x* – *x*_*i*_)^2^ + (*y* – *y*_*i*_)^2^, *R*_0_ is the initial packet radius, *t*_*i*_ is the time since the release of this particular packet, *D* is a diffusivity that governs the packet growth, meant to account for molecular diffusivity and the effects of small eddies and *C*_0_ sets the initial concentration amplitude. The total odor(*x, y, t*) is then the sum over all packets that have been released up to time *t*. The packet center was computed the following way,

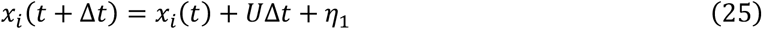

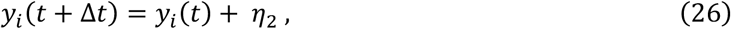

where *U* denotes the mean wind velocity and *η*_1_ and *η*_2_ are Gaussian white-noise perturbations with mean 0 and standard deviation 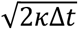, representing the effects of turbulent dispersion with eddy diffusivity κ.

In general, parameters were chosen to be physically realistic and also give concentration time-series and odor plumes that were qualitatively similar to those in the videos. To set *C*_0_, we defined the detection threshold to be 1 and enforced that an agent more than 1.6 standard deviations away from an initial packet would not be able to detect its presence. See Table 2 below.

**Table 1:** List of parameters used for odor plume simulations

| Parameter | Explanation | Value |
| --- | --- | --- |
| $U$ | Wind speed | 36mm/s – 300mm/s |
| $D$ | Packet growth diffusivity | 10mm<sup>2</sup>/s – 52mm<sup>2</sup>/s |
| $\kappa$ | Eddy diffusivity | 1000mm<sup>2</sup>/s |
| $\lambda$ | Packet release rate | 5Hz |
| $R_0$ | Initial packet radius | 10mm |
| $C_0$ | Initial packet intensity | 3827.24 (a. u.) |
| $K$ | Odor detection threshold | 1 (a. u.) |

The order of magnitude for *D* was set by the fact that attractive odorants for *Drosophila melanogaster* tend to have molecular diffusivities of around 10mm^2^/s, eg. Ethyl acetate. The eddy diffusivity κ was set in accordance with (Drivas et al., 1996). The release rate and initial size were chosen to be similar to those in (Farrell et al., 2002). The wind speed was chosen to be similar to those used experimentally in (Demir et al., 2020) and (Alvarez-Salvado et al., 2018).

Additionally, to improve computational efficiency, packets were no longer tracked once their *x* position was so large that even if all released packets were at that position, the sum of their contributions would still be less than the detection threshold.

10,000 agents were initialized with uniformly distributed initial position and angle, with *x* between 50mm and 400mm, *y* between -110mm and 110mm and 0° < *θ* < 360°, where *x* and *y* positions are defined relative to the source location, as in Figure 3. Plumes were simulated for enough time steps so that the expected *x* position of a packet released at time 0 would be equal to the maximum initial *x* for navigating agents, before navigating agents were introduced and simulated for 120s. Once again, a trajectory’s success was defined by whether it got within 15mm of the source location.

To define the antenna-sensing region, space was discretized into “pixels” with 0.154mm as the pixel width, matching the spatial resolution of the high frequency plume. The concentration was then computed by averaging over the pixels in an elliptical region, with the region defined as in the previous section.

To set the level of noise added to the *I* and *F* filters, we first computed a characteristic *I* value in the simulated high intermittency plume, *I*_0_, by averaging *I* values over a region 192mm < *x* < 205mm and 0mm < *y* < 9mm and then averaging over the length of the simulation. We did the same for *F* values in the simulated high frequency plume to obtain *F*_0_. The values we obtained were *I*_0_ = 0.776 and *F*_0_ = 3.14. We then used 5% of these values as the standard deviation for Gaussian white noise to be added to the output of the *I* and *F* filters respectively at each time step. We also used *I*_0_ and *F*_0_ as representative *I* and *F* values in order to assign a single relative filter weight with which to color each set of gains in Figure 4G.

---

**Figure 4-figure supplement 1.**
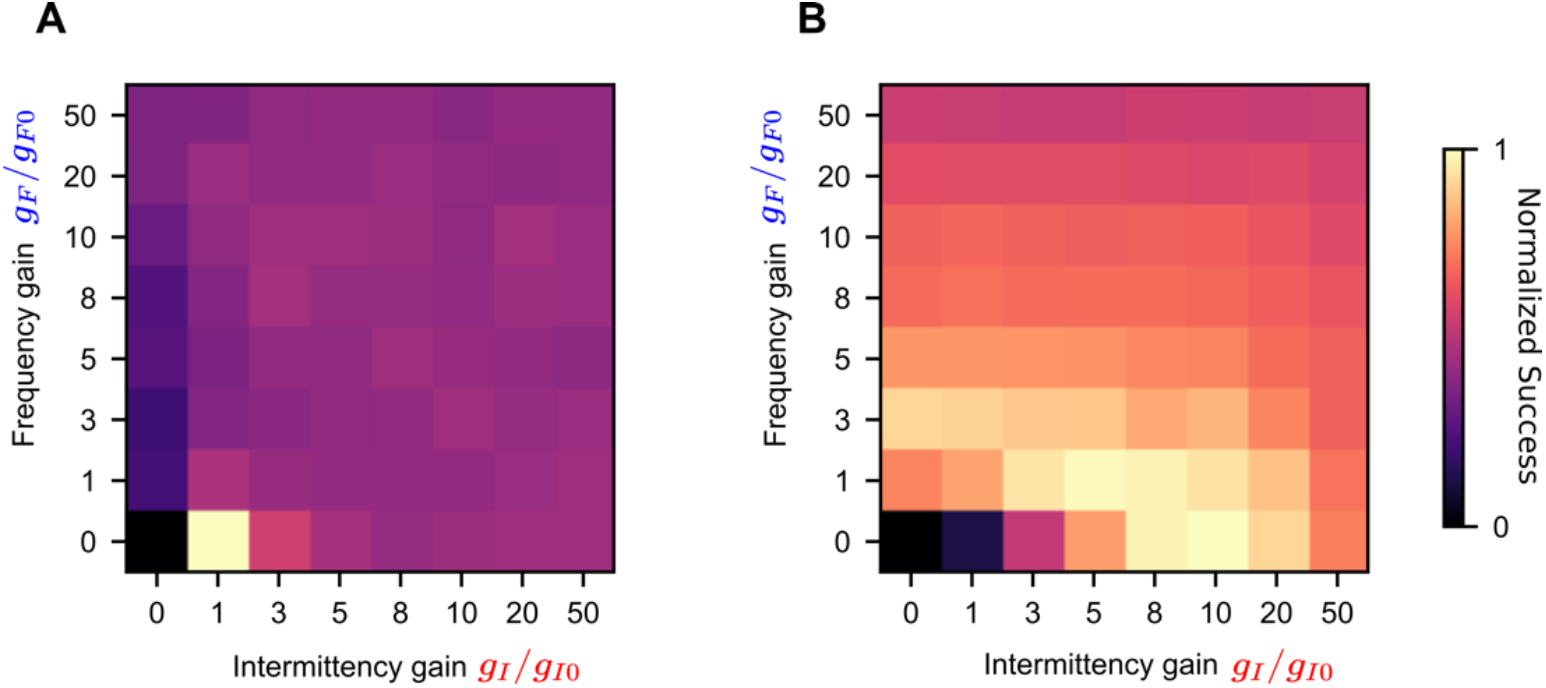
Performance for different sets of gains without filter noise: Normalized success in the simulated high intermittency plume **(A)** and simulated high frequency plume **(B)** for different sets of *I* and *F* gains. We see strong performance in B for a large region at high intermittency gains *g*_*I*_.

## Notes

### Competing Interest Statement

The authors have declared no competing interest.

